# SVJAM: Joint Analysis of Structural Variants Using Linked Read Sequencing Data

**DOI:** 10.1101/2021.11.02.467006

**Authors:** Mustafa Hakan Gunturkun, Flavia Villani, Vincenza Colonna, David Ashbrook, Robert W. Williams, Hao Chen

## Abstract

Linked-read whole genome sequencing methods, such as the 10x Chromium, attach a unique molecular barcode to each high molecular weight DNA molecule. The samples are then sequenced using short-read technology. During analysis, sequence reads sharing the same barcode are aligned to adjacent genomic locations. The pattern of barcode sharing between genomic regions allows the discovery of large structural variants (SVs) in the range of 1 Kb to a few Mb. Most SV calling methods for these data, such as LongRanger, analyze one sample at a time and often produces inconsistent results for the same genomic location across multiple samples. We developed a method, SVJAM, for joint calling of SVs, using data from 152 members of the BXD family of recombinant inbred strains of mice. Our method first collects candidate SV regions from single sample analysis, such as those produced by LongRanger. We then retrieve barcode overlapping data from all samples for each region. These data are organized as a high dimensional matrix. The dimension of this matrix is then reduced using principal component analysis. Samples projected onto a two dimensional space formed by the first two principal components forms two or three clusters based on their genotype, representing the reference, alternative, or heterozygotic alleles. We developed a novel distance measure for hierarchical clustering and rotating the axes to find the optimal clustering results. We also developed an algorithm to decide whether the pattern of sample distribution is best fitted with one, two, or three genotypes. For each sample, we calculate its membership score for each genotype. We compared results produced by SVJAM with LongRanger and few methods that rely on PacBio or Oxford Nanopore data. In a comparison of SVJAM with SV detected using long-read sequencing data for the DBA/2J strain, we found that our results recovered many SVs missed by LongRanger. We also found many SVs called by LongRanger were assigned with an incorrect SV type. Our algorithm also consistently identified heterozygotic regions.

## 1 Introduction

Linked-read whole genome sequencing (WGS) methods, such as the 10x Chromium platform, attach a unique molecular barcode to each high molecular weight DNA molecule. The samples are then sequenced using short-read technology, such as the Illumina platform. During analysis, reads sharing the same barcode are aligned to adjacent genomic locations. Linked-read sequencing provides a inter-mediate platform between short read sequencing and single molecule long-read technology (e.g. PacBio and Nanopore platforms). Because DNA are sequenced using short read technology, linked-read data has high base-level quality and is inexpensive to obtain. Further, the barcode unique to high molecular weight DNA allows the detection of large structural variants (SV), roughly in the range of 1 Kb – 2 Mb, which are difficult to detect using short-read sequencing.

After aligning the sequencing reads to a reference genome, the molecular barcodes can be visualized in a two dimensional space, with each axis presenting a genomic location (i.e., matrix view). When both axis represent the same genomic region, the number of barcodes shared by any location are concentrated on the diagonal line. Aberrant patterns in these image indicate the presence of SVs. Figure 1 shows a 351 kb region of Chromosome (Chr) 4 for three samples. Figure 1a belongs to B6 mouse (reference) and Figure 1b belongs to D2 mouse (alternative). The sample in Figure 1c shows a heterozygous deletion.

**Figure 1:**
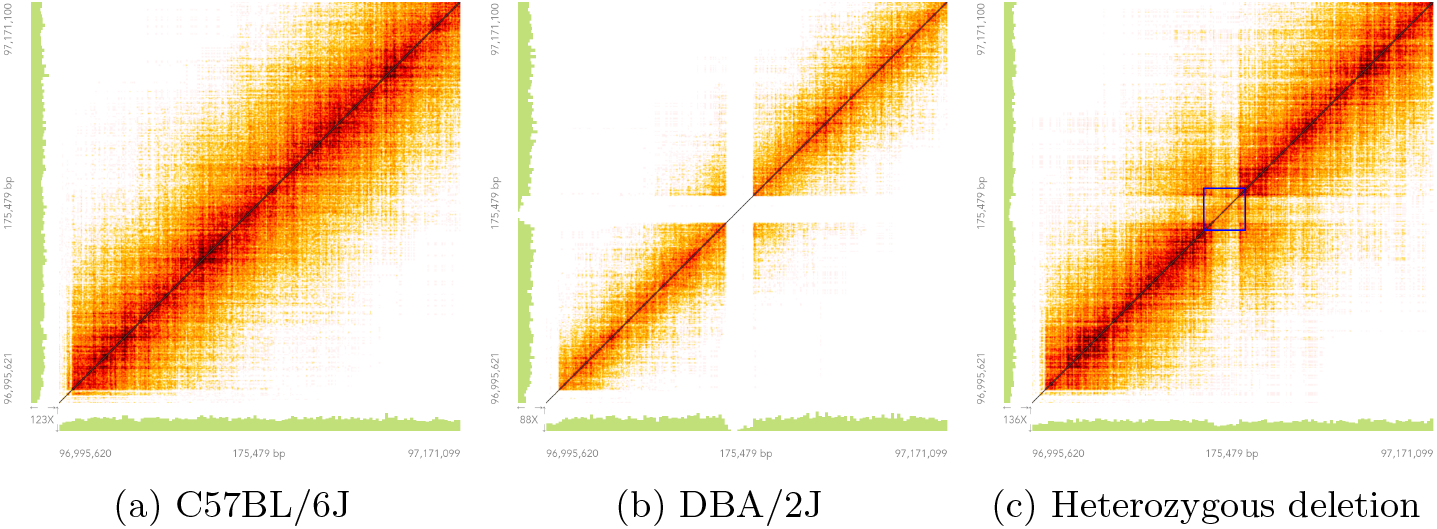
A 175 kb region on Chromosome 4 showing the reference allele (a), a homozytotic deletion (b) and a heterozytotic deletion (c)

Most SV calling methods for the linked read data, such as LongRanger, analyze one sample at a time. We have found that the reported SVs for a given location were often inconsistent across multiple samples. We developed a method, SVJAM (structural variant joint analysis by machine learning), that detects and genotypes large structural variants (SVs) from linked-read whole genome sequence data generated on the 10x Chromium platform. We tested SVJAM on a data set containing 152 BXD recombinant inbred mice. The BXD family of mice were generated by crossing female C57BL/6J (B6) to male DBA/2J (D2) mice, followed by generations of inbreeding of the offspring. Recently, these strains have been fully sequenced using linked-read technology (Ashbrook et al., 2021). By analyzing 152 individuals jointly, SVJAM produces consistent and accurate SV detection and genotyping. In addition, SVJAM also detects heterozygotic SVs with high accuracy.

## 2 Methods

### 2.1 Overview

An overview of our method is presented in Figure 2. Like other joint analysis methods for genetic variants (Yun et al., 2021), SVJAM takes candidate regions suggested by other methods that analyzes one sample at a time. The input data for SVJAM are barcode overlapping images for these candidate regions from all samples. These data are generated by using the Loupe application of the 10x Chromium Platform in the form of ”matrix view” (Figure 1). Although not open source, the Loupe browser is freely available from 10x Genomic website ^1^. The pixel intensity of these images corresponded to the depth of barcode overlapping between two genomic locations defined by the x- and y-axis. SVJAM was developed using a data set containing linked-read WGS of 152 BXD family of recombinant inbred (RI) mice (Ashbrook et al., 2021), primarily using 2,114 candidate regions on Chr 1 suggested by LongRanger ^2^ (321,328 matrix view images).

**Figure 2:**
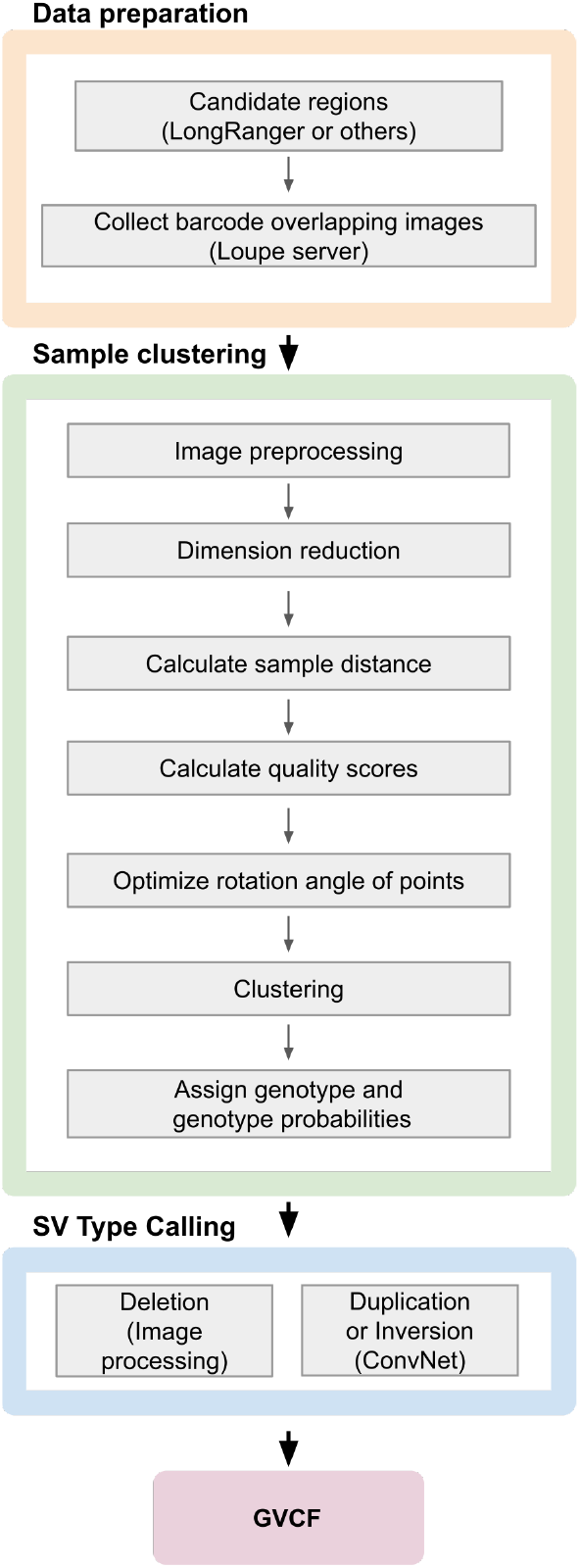
Workflow of SVJAM

After obtaining these images, we ran the pipeline for each genomic location. First, we determine the presence of an SV. For deletions, we use image processing techniques to analyze the lack of barcode overlapping (i.e., white pixels) along the diagonal of the images, which indicates deletions. We trained a convolutional neural network (CNN) for the detection of duplication and inversions. CNN is a successful supervised machine learning technique for object detection. We trained the model by using 12,000 images that belong to regions in which at least 40 samples were annotated as having duplication or inversion. We labelled the training set automatically by using a clustering algorithm. Then, we trained the model by using a network containing 18 convolution, max pooling, dropout and flatten layers.

The clustering algorithm is applied to each genomic region and begins by preprocessing the images. We organize the images as columns of a big matrix and apply dimension reduction techniques to visualize the representations of individuals in 2-dimensional plane. The ranges of the axes after principal components analysis (PCA) is usually not symmetric which may bias the clustering methods. To overcome this, we generated a custom distance matrix which is not symmetric in terms of *x* and *y*. Some projections may also need to be rotated to be ready for clustering. The coefficient in the new distance metric and the rotation angle is determined by optimizing a quality function. The number of clusters represent the number of genotypes. A third cluster exists when there are samples with heterozygous SVs present in a region. We developed a formula to determine the number of clusters. After clustering, we assign a vector to each individual that shows the probability of it to be a member of each cluster. The output of the pipeline is a text file in the Genomic Variant Call Format (GVCF) that contains general information about each SV and genotype, as well as quality of the call for each individual.

### 2.2 Data preprocessing

We converted matrix view images of each individual from RGB to grayscale using the ITU-R 601-2 luma transformation. This does not effect the processing since the pixels of the images does not lose color intensity information. The images are cropped to remove the parts consisting the read depth and the numbers about the genomic location and base pair information. There are 152, 1018 × 1018 pixel images for each genomic location. After flattening the representation matrices of these images, we arranged each sample as a column vector to have a 1 million times 152 dimensional data frame. To sum up, the last matrix represents the gray scale images of one particular genomic region of all the individuals, each individual in one column.

### 2.3 Dimension reduction

After standardizing the features by removing the mean and dividing by the standard deviation, we applied principal component analysis (PCA) to project the data into two dimensional plane. Figure 3 shows the results of PCA applied to a 760 kb region on Chr 17. The axes of the figure are the first two principal components of PCA. Each red point in the figure represents one individual.

**Figure 3:**
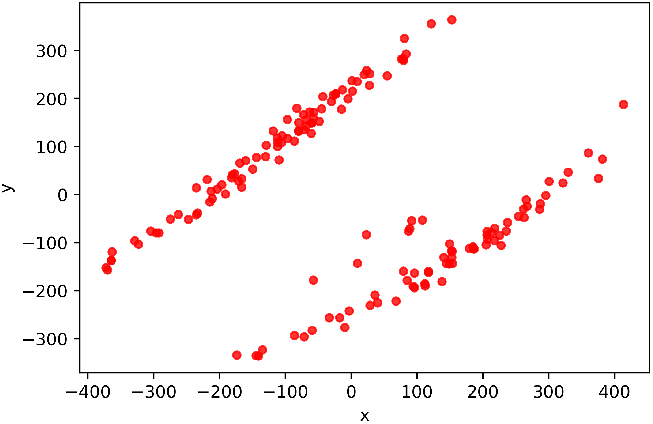
Scatter plot of the first two principal components of PCA results of 153 samples, 760 kb region on Chr 17.

### 2.4 Clustering and metrics

The consistency of the structural variants among the samples makes unsupervised machine learning approaches appropriate for the joint analysis of SVs. After the preprocessing and dimension reduction, we formed an algorithm including hierarchical clustering with some modifications. To merge two sets, we used single linkage criterion which combines the clusters by using the minimum of the distances between all elements of the two sets.

The default distance metric in these algorithms is Euclidean metric, yet we could not use that since the ranges of two principal components are different. To get a useful distance metric we broke the symmetry of the weights in horizontal and vertical distances. We defined a weighted Manhattan distance (weighted taxicab metric or weighted *L*^1^ − *norm*) between the points *p* = (*p*_1_, *p*_2_) and *q* = (*q*_1_, *q*_2_) in the plane as follows:

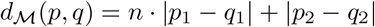

It is straightforward to show that this is a metric function hence this defines a topology on ℝ^2^ for a given *n* ∈ ℝ. For the values of *n* greater than 1, this function puts a weight to horizontal direction which would work as if a punishment in the clustering algorithm.

For each genomic region, we formed a distance matrix *M*_*ij*_ = *d*(*s*_*i*_, *s*_*j*_) where *s*_*i*_ = (*s*_*i*1_, *s*_*i*2_) and *s*_*j*_ = (*s*_*j*1_, *s*_*j*2_) are the coordinates of the first two principal components of *i*^*th*^ and *j*^*th*^ samples, respectively. We use this matrix for all the distance calculations, so that clustering algorithms on a region is based on this custom metric.

The best weight *n* in the distance function is automatically determined by the machine for each genomic region. To this end, we need to define some performance metrics for clustering. We also use these metrics to determine the number of clusters. We will present the explanations of these in the next section. Before that, we present one more arrangement.

The PCA results in some genomic locations needs to be rotated since they have an oblique tendency. After a certain rotation, they become ready for clustering. We use the following planar rotation matrix *R* to rotate the points.

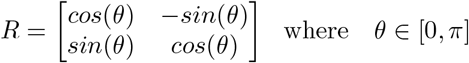

The rotation angle is also determined by the machine by using the performance metrics defined in the next section.

#### 2.4.1 Performance scores

We calculated a score for measuring the quality of clustering. First set of formulas below are to see how well the elements within the clusters gathered. The total distance of an element *x* to all other elements in the same cluster is called *wd*(*x*) for “within distance of x”. This quantity is divided to the number of elements in that cluster to get “within mean of x”, shortly *wm*(*x*). Then, we got “within mean” by finding the sum of within distance of each element and dividing it to the total number of elements.

The formulations are as follows. Let *d*_ℳ_ be the weighted Manhattan distance function that we defined above, *C*_*x*_ be the set of elements that are in the same cluster with *x, C* be the set of all elements and |*C*| be the cardinality of C. Then we find *wm* as:

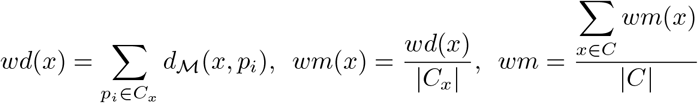

We implemented similar calculations for the elements between different clusters to see the quality of the separation of the clusters. The first equation finds the total distance of an element *x* with the elements in other clusters. Second equation aims to find the mean distance of *x* with elements in other clusters by dividing the total distance by the number of elements in clusters not including *x*. The last equation finds the sum of all distances with elements in different clusters and divides it to the total number of elements to find the mean.

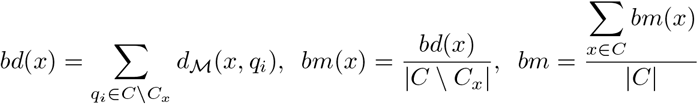

where *bd*(*x*), *bm*(*x*) and *bm* stands for ”between distance of x”, ”between mean of x” and ”between mean”, respectively.

By using these identities, we assigned a quality score for each clustering. We set the cluster quality as follows:

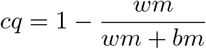

This gives a comparable scoring between different clustering cases. In an ideal clustering with 3 clusters, the sets would be apart from each other as much as possible whereas the points in each set are close to each other. Then, *wm* will be very small but *bm* will be large to make *qs* close to 1. The opposite case would make *qs* close to 0.

Besides putting a measure for overall quality of clustering, *qs* is also used to determine the rotation angle that we discussed in the previous section. The best angle for the rotation of the PCA plot of a genomic location is the one that makes the quality score highest. For instance for Figure 3, the best angle is 45°. The PCA result for the same location with 45° rotation is shown below.

We also used the identities above to determine the best weight *n* for the distance function *d*_ℳ_(*p, q*) = *n* · |*p*_1_ − *q*_1_| + |*p*_2_ − *q*_2_| of two points *p* = (*p*_1_, *p*_2_) and *q* = (*q*_1_, *q*_2_). Notice that, in this equation *d*_*ℳ*_(*p, q*) increases if *n* increases, which leads to an increase in *wm* and *bm* that are defined above. And, if the other variables remain constant, *qs* increases as well. Therefore we could not use *qs* as it is, to compare the clustering for different values of *n*. Instead, we used the lag between *wm* for two consecutive values of *n*. This gave a good measure for the improvement in the clustering when we increase *n*. We put a threshold to remove the irrelevant changes and we chose *n* that results in the best improvement for each genomic location.

#### 2.4.2 Number of Clusters

Determining the best number of clusters in unsupervised learning practices is usually not an easy task. In our case, for each genomic location, three types of clustering are possible as shown in Figure 5.

**Figure 4:**
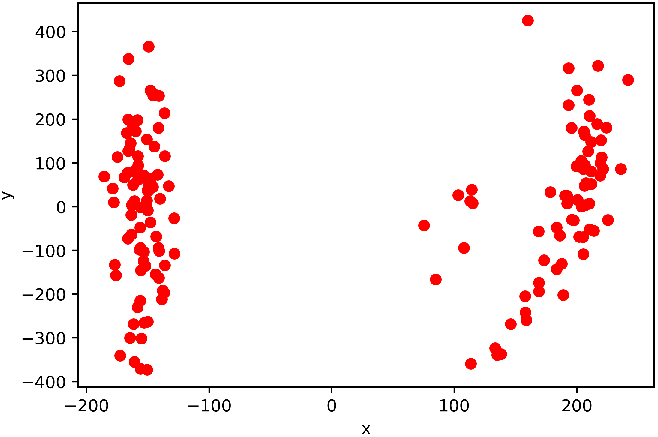
Scatter plot of the first two principal components of PCA results of 153 samples with 45° rotation, a 760 kb region on Chromosome 17

**Figure 5:**
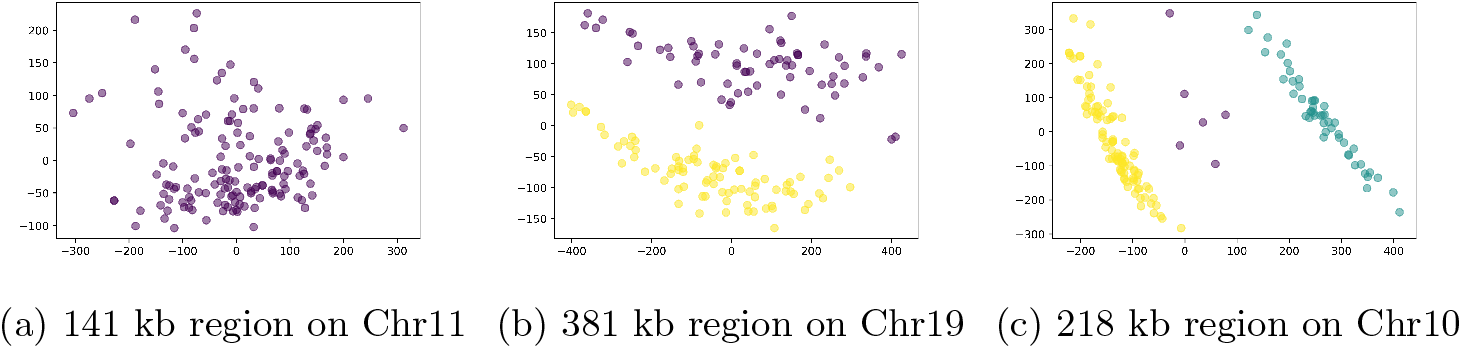
Types of clusters

There may be an assembly error or all the samples may have the same variant. Both cases result in the samples to be in one cluster (Figure 5a). There are two clusters if there are homozygous variants in some samples and alternative variants in the others whereas there are no heterozygous variants (Figure 5b). Three clusters are formed if there are both homozygous and heterozygous variants (Figure 5c). To separate the first case from the other two, we used Calinski-Habarasz score (CH-score) (Calinski and Harabasz (1974)). We put a threshold to distinguish one cluster cases from the others. We could not use CH-score and other known clustering metrics to distinguish the other two types. One reason for that is they are by default using the Euclidean distance, hence we needed to change it to a custom metric as explained above. The other reason is that our data points need a refined measure rather more general and rough measures.

To find the best number of clusters (either 2 or 3), we first find the centroids of the clusters. In the following equation 𝒞_*i*_ is the centroids for the homozygous reference (*i* : *ref*) and homozygous alternative (*i* : *alt*)

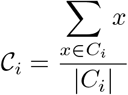

where *C*_*i*_ is the corresponding cluster, *x* is the planar coordinates of the representation of an individual in *i*^*th*^ cluster, |*C*_*i*_| is the cardinality of *C*_*i*_ and the sum on the numerator is calculated coordinate-wise.

In this step, we assume that there are 3 clusters, then either we accept or reject that by using following calculations. If it is rejected, then the pipeline proceeds with 2 clusters, otherwise it proceeds with 3 clusters. We calculate the normalized mean distances between the heterozygous individuals to the centroids C_*ref*_, C_*alt*_ of other clusters.

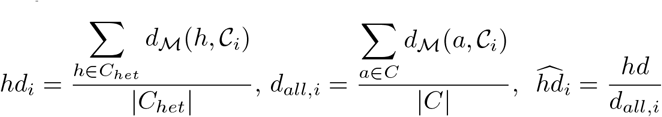

The first sum is taken over the individuals that belong to the heterozygous cluster. The summand is the distance of a heterozygous individual to 𝒞_*i*_, *i* ∈ {*ref, alt*}. The denumerator is the number of heterozygotic individuals. The second sum finds the mean distance of all individuals to centroids of the clusters where *C* is the set of all individuals. Lastly, 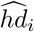 is the normalization of *hd*_*i*_. We reject the assumption of having 3 clusters if the ratio between *hd*_*ref*_ and *hd*_*alt*_ is bounded by a treshold from below and above.

To sum up, if there is enough distance (normalized) between the heterozygous cluster to both homozygous clusters, then we accept that there are 3 clusters. Else, if the heterozygous cluster is close enough to one the other 2 clusters, we reject the hypothesis of having 3 clusters.

#### 2.4.3 Membership Probabilities

The last set of calculations assigns a membership probability vector to each individual, that is the probabilities of an individual to belong each cluster. Unlike the scores that measures clustering performance, membership probabilities measure the affinity to the parental strains. We first calculated the total distance of all elements to the reference strain, then divide it to the total number of individuals to get the mean distance to the reference strain, call it 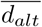. Similarly we calculated 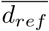 to get the mean distance to the alternative strain.

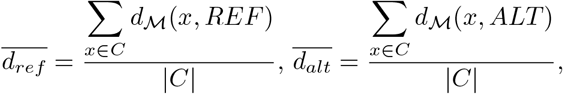

where *REF* and *ALT* are the coordinates of the reference and alternative strains. Then, the normalized distance of an individual to the reference and alternative strains are calculated as:

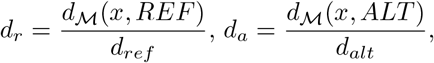

The figure below is the (*d*_*r*_, *d*_*a*_) pairs in the coordinate plane having distance of the individuals to reference and alternative strains as *x* and *y* axes, respectively.

To determine the distance of an individual to being heterozygous, we take the distance of (*d*_*a*_, *d*_*r*_) pair to the diagonal line (*y* − *x* = 0) of the above figure. So, we get

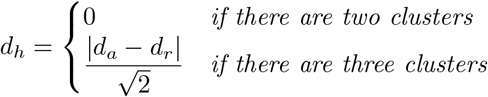

The last part is calculating the membership probabilities by using these distances. We explain the probability formulas on this sample case.

We will calculate the probabilities of the highlighted point (*y*) in Figure 7 to be a member of Green cluster (includes reference), Purple cluster (includes alternative) and Yellow cluster (heterozygous individuals).

**Figure 6:**
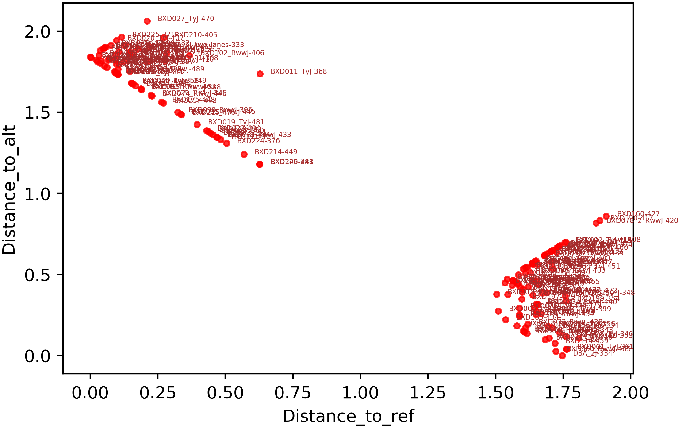
Scatter plot of (*d*_*r*_, *d*_*a*_) coordinates of 152 samples of a 760 kb region on Chromosome 17

**Figure 7:**
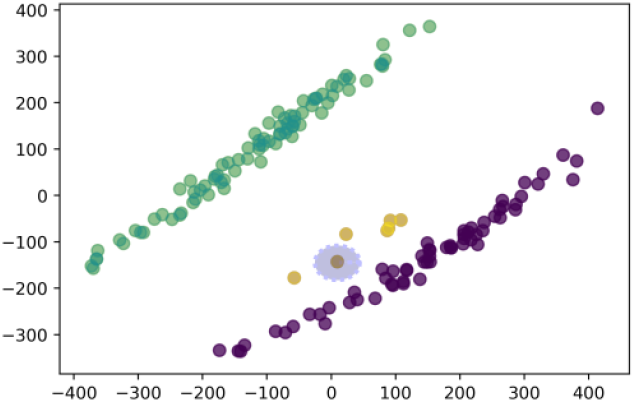
Scatter plot of the first two principal components of PCA results of 153 samples, 760 kb region on Chromosome 17.

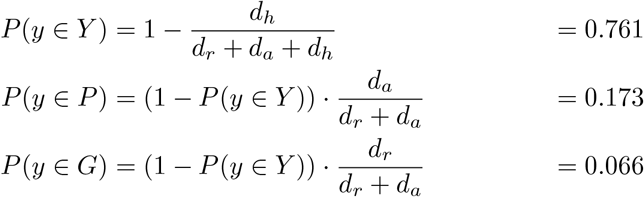

The vector of predicted probabilities of the highlighted individual *y* to be a member of the clusters that include the samples with homozygous reference, heterozygous and homozygous alternate variants is (0.066, 0.761, 0.173).

The membership probabilities for the individuals in other clusters are calculated similarly, the only change is the order of the coordinates of vector (*d*_*h*_, *d*_*r*_, *d*_*a*_) in the formulas.

### 2.5 Detecting Structural Variants

We determine the type of SVs in each candidate genomic region once the clustering step was finished.

#### 2.5.1 Deletions

We used image processing techniques to detect the presence of deletions, which were manifested as the lack of barcode overlap along the diagonal line in the matrix view images, with the appearance of a white cross (Figure 8a). These white strips were detected by an algorithm that are tolerant to a small percentage of grey pixels. After analyzing these we report the beginning and ending of the deletions and their approximate lengths.

**Figure 8:**
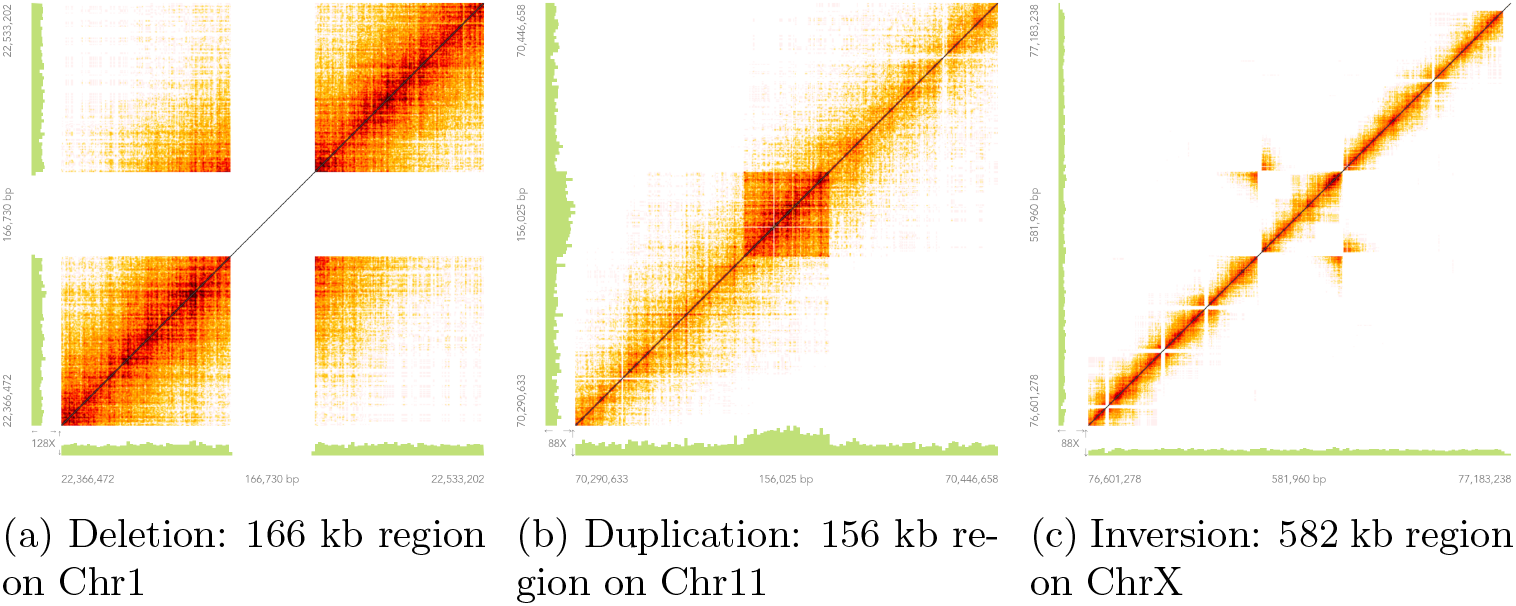
Sample images for deletion, duplication and inversion

#### 2.5.2 Duplication and Inversion

We trained a CNN to classify the images that include duplication (Figure 8b) or inversion (Figure 8c). A dataset was arranged for training and validation having 75 regions, totally 11,400 images. These images were selected from regions where LongRanger reported at least 40 samples having duplications or inversions. These locations are genotyped through the joint calling algorithm and we got the labelling as an output. Hence, the labelling of the images that would be used for supervised learning is set by the unsupervised algorithm rather than a manual work. Then the data set was split as 70% and 30% randomly into training and validation sets, respectively. Training accuracy is calculated on the validation set after each epoch, and the best weights that gives the highest accuracy was chosen for the model. We selected another set of 75 regions for testing the model manually. The images are compressed to 512×512 pixels for training and CNN is performed by using NVIDIA GeForce RTX 3090 GPU.

We used Keras API on Tensorflow framework. The model consists of 18 convolution, max pooling, dropout and flatten layers. The loss function that the model tries to minimize is categorical cross entropy and the accuracy is measured by categorical accuracy. Adam optimizer function is used to update the parameters with a learning rate 0.0001. The batch size is 64 which means 64 images are processed in one gradient update. The model has a categorical accuracy of 0.9983 on validation set. These weights are used to predict duplication and inversion in over 4 million images.

### 2.6 Genotyping each sample and producing output

The presence of SV in each genomic region is first determined by the number of clusters. Regions with one cluster were discarded in our current implementation because the B6 mice, where the reference genome was based on, was one of the samples in our analysis. However, this behavior can be easily modified. For regions with two or three clusters, each sample were assigned the genotype of the majority of the cluster. This approach corrected the occasional mistakes made by the CNN in determining inversions or duplications.

The location of the deletion was determined by checking the beginning and ending of white strips in the images. If the strip is less than 1000 bp, we skipped that in case the image might be noisy. After we verify the existence of duplications and inversions, we rely on the positions annotated by LongRanger, even it finds them in only a few samples.

After SVJAM detected and genotyped the SVs, we collected all the information in a text file in the Genomic Variant Call Format (GVCF). The file includes clustering quality scores for each region and membership probability vectors for each individual besides the common information as beginning and ending positions, sizes, etc. We also include CIPOS and CIEND to the GVCF file that shows the confidence interval for POS and END. So, CIPOS and CIEND are intervals specifying the region of uncertainty around the POS and END values.

### 2.7 Long-read genome sequencing of the DBA/2J mice and data analysis

Two healthy adult male DBA/2J mice from the colony at the University of Tennessee Health Science Center were used. Spleen was used for DNA extraction. Oxford Nanopore (ONT) sequencing were conducted by using a Promethion instrument by DNA Link (Los Angeles, CA 90015). A total of 779,223 reads were obtained (14.25 billion bases, N50 of 29,205). Pacific Biosciences (PacBio) HiFi data were generated by the DNA sequencing core facility at the University of Wisconsin. A total of 2,674,984 reads were obtained(28,568,875,629 bp, N50 of 11,307 and largest contig of 44,747). All sequencing runs were conducted using manufacturer suggested protocols

ONT data were mapped to the reference genome mm10 using minimap2 (version 2-2.17) (Li, 2018). SVs were detected using Sniffles (version 1.0.12) (Sedlazeck et al., 2018), SVIM (version 1.4.2) (Heller and Vingron, 2019), and NanoVar (version 1.3.9) (Tham et al., 2020). PacBio data were mapped to mm10 using pbmm2 ^3^. SV were detected using Sniffles.

## 3 Results

### 3.1 Dimension reduction

We applied various linear and non-linear dimension reduction techniques. Three of them are shown in Figure 9 for a 243kb region in Chr 4. First figure shows the result for principal components analysis, second and third figures shows the embedding for non-linear techniques, t-distributed Stochastic Neighbor Embedding (t-SNE) and spectral embedding. We get the best embedding results by using PCA, hence we decided to use it for the projection for all the regions.

**Figure 9:**
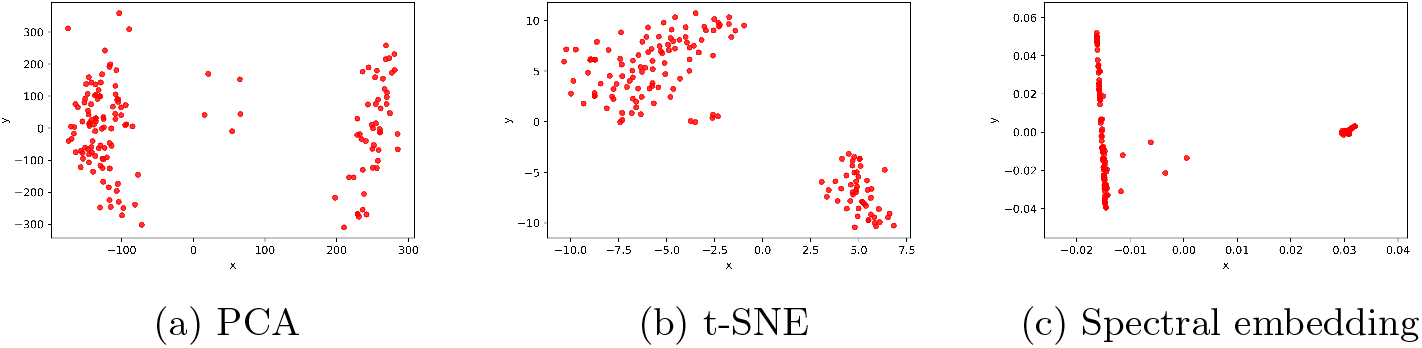
Dimension reduction examples

### 3.2 Distance metric

After we embed the data into 2-dimensions by PCA, we applied different clustering methods as k-means and hierarchical clustering. The result of hierarchical and k-means clustering for a 243kb region in Chr 4 is shown in Figure 10.

**Figure 10:**
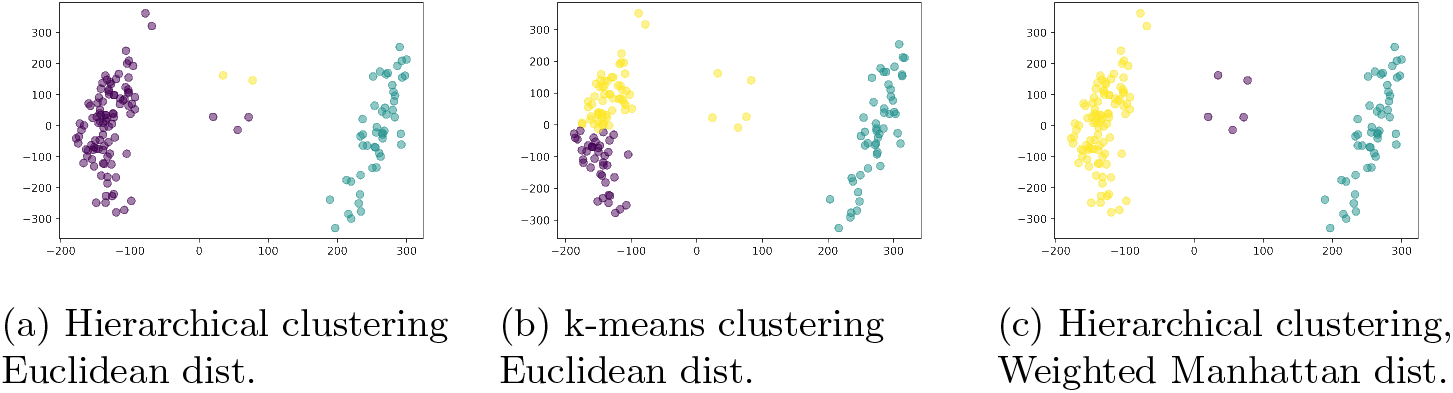
Dimension reduction examples

The first two images are formed by using Euclidean metric (*L*^2^ −*norm*) which is the default metric distance for most machine learning libraries as Scikit-learn for Python. Clustering by using the Manhattan distance (*L*^1^ − *norm*) produced the same results. We expect to have 3 clusters in this region as having two big clusters on the left-right and one small cluster with 5 points in the middle as in Figure 10c. However, since the embedding after PCA has different *x* and *y* variance and these norms are symmetric in terms of *x* and *y*, we get these unexpected figures. To solve this issue, we introduced a new metric that is weighted Manhattan distance (*d*_*ℳ*_) which gives the desired clustering for all the regions. Figure 10c is the result of hierarchical clustering using weighted Manhattan distance with *n* = 5 in the formula. The metric *d*_*ℳ*_ is explained in detail in Methods section.

### 3.3 Rotating the points

The data points for some regions are structured as in Figure 11a after PCA embedding. The points are rotated to improve clustering results. The rotation results with different angles are shown in Figure 11b-11f. The best angle of rotation for this example is 50°, in general the angle is determined by the quality scores for each region which is explained in Methods section.

**Figure 11:**
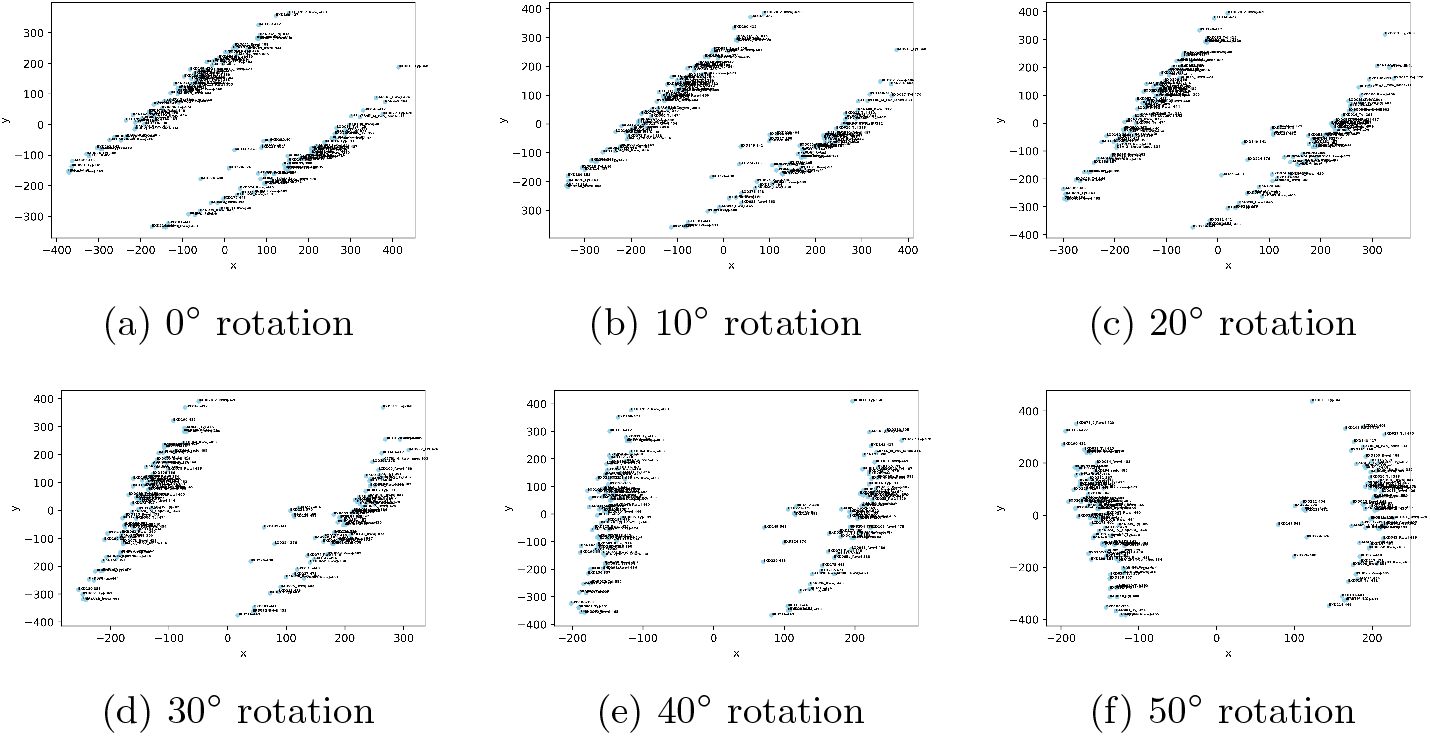
Rotation of the points

### 3.4 Analysis of SVs on Chr 1 by SVJAM

We jointly analyzed Chromosome 1 of 152 whole genome sequences of BXD family and ran SVJAM for 2,114 regions including 321,328 individual images. SVJAM found that 328 of these regions contain large SVs, 191 of them were deletions, 62 of them were duplications and 75 of them were inversions. The minimum size of SV was 1002 bp, and the mean of the sizes of all SVs was 33,480 bp. The summary statistics for sizes was shown in Table 1.

**Table 1:** Summary statistics for the size of structural variants

| Min. | 1st Qu. | Median | Mean | 3rd Qu. | Max |
| --- | --- | --- | --- | --- | --- |
| 1002 | 4226 | 9394 | 33480 | 39767 | 1051306 |

We compared the SVs of the D2 mice reported by SVJAM with the original SVs reported by LongRanger, and three result sets reported using different tools on the D2 ONT data, and one result set produced using PacBio data. We focused the analysis on the largest chromosome, Chr1. The distribution of SV size for different methods was show in Figure 12. SVs reported by LongRanger were the largest, centered around 38 kb. SVJAM retained some of these SVs but also increased the proportion of SVs at about 9 kb.

**Figure 12:**
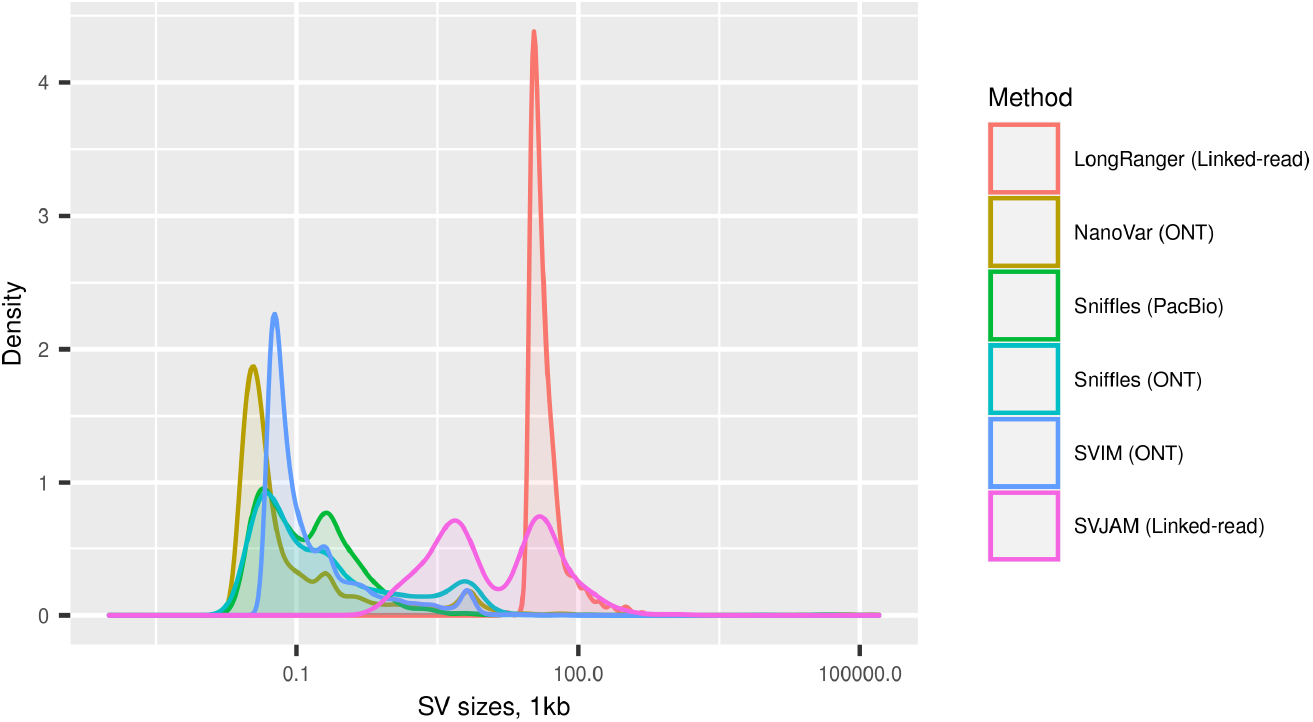
Size distribution of structural variants identified by different methods

The total number of SVs and the number of overlapping regions were shown in Figure 13. A total of 185 SVs were only found by LongRanger which were most likely false positives. All the SVs detected by SVJAM were reported by at least one other method based on long read sequencing.

**Figure 13:**
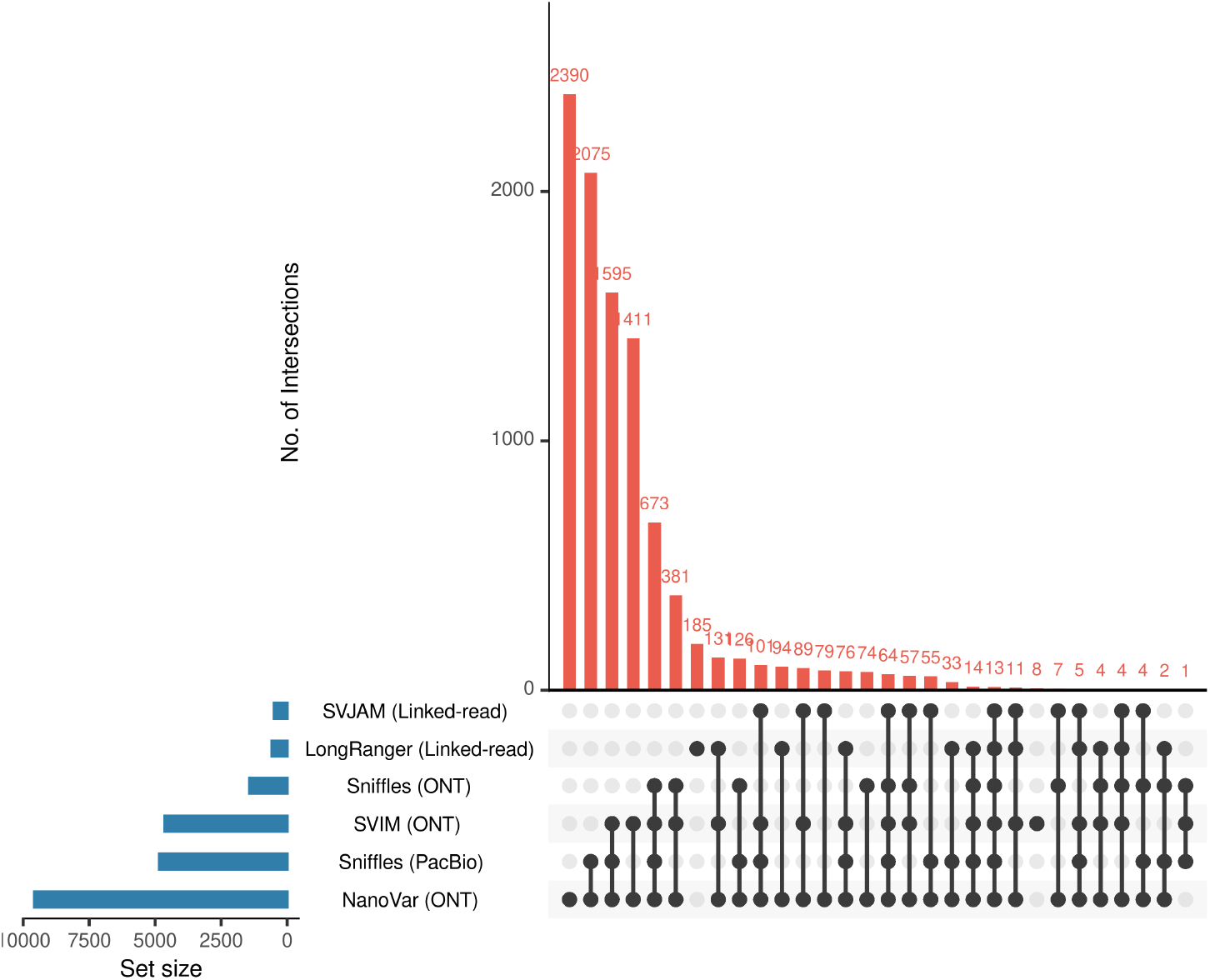
Number of overlapping regions for Chromosome 1 analyzed by using different SV callers. The blue bars on the left showed the number of SVs. The black bars showed the intersecting subset of methods and the orange bars showed the number of elements in those subsets.

## 4 Discussion

We developed a method, SVJAM, to jointly detect and genotype several types of structural variants, including deletion, inversion, and duplication, from linked-read WGS data. SVJAM used imaging processing techniques to detect deletion and used CNN to detect duplication and inversion. The genotype of each sample was determined by a clustering method. We compared the SVs detected by SVJAM on Chr 1 for the D2 mice with those reported using the same data by LongRanger, and those reported by four other methods using long-read PacBio or ONT data. All variants reported by SVJAM were corroborated by at least one other method, while LongRanger reported many SVs not supported by the long read data. Furthermore, joint calling makes it possible to identify heterozygotic SVs accurately.

Similar to most methods of joint analysis of genomic variants, such as GLNexus (Yun et al., 2021), SVJAM does not detect SV by default. Instead, it takes candidate regions suggested by other methods to start the joint analysis. We used LongRanger to obtain these candidate regions. We required the candidate regions to have at least 5 individuals with SVs. We observe that LongRanger often produces inconsistent results, in both calling variants and determining the type of SV. Hence, we apply SVJAM for all the candidate regions regardless of the type of SV reported by LongRanger. Although we used LongRanger to suggest candidate regions, SVJAM can work on any arbitrary genomic region.

Supervised and unsupervised learning practices have unique advantages and disadvantages. In this project, we try to compensate a drawback of one by using a benefit of the other. For example, labelling the training data set in CNN is a time and effort consuming progress. We used clustering algorithm to label the images in each region. The individuals were labelled by checking that whether they were in the same cluster with the homozygous reference, homozygous alternative or they belong to a third cluster, heterozygous samples. By this way we were able to collect 12,000 labeled images with minimal effort. Another example was to use clustering to correct the few mistakes made by the CNN in the genotype stage. We first used the CNN to call duplication or inversion on each sample. We then assign the cluster to the majority of the SV type. If there were samples having SV type predicted by the CNN that were not the same as that of the cluster it belong to, we consider those mistakes made by the CNN and reassigned the SV type of those samples based on their cluster identity.

The formulations in the clustering algorithm did not depend on the data. Hence, we believe that they can be applied to other problems as well. The clustering quality score formula was used to evaluate the performance of the clustering independent from the method and the data. A formula based on the distances of individuals to the centroids of the clusters was used for determining the number of clusters. Lastly, the membership probability vectors assign each point a vector that estimates the belonging of the point to each cluster. These could easily be generalized to problems that include more number of clusters.

There were also many limitations in SVJAM. Similar to LongRanger, SVJAM does not report insertions. This is because insertions do not have clear pattern in Matrix View images. A second limitation, also because we used SV candidates suggested by LongRanger, is that the result set lacks sensitivity to SVs smaller than 500 bp. Another limitation is that the location of SV are either determined by image processing (for deletion) or taken the average reported by LongRanger, and thus are not exact. A third limitation is that the analysis is conducted using image data retrieved from the Loupe browser, rather than numeric data. The distinctive pattern of each SV in matrix view images facilitate the algorithm development process. We also tested obtaining the same data in csv format from the Loupe browser and found the time saving to be minimal. We therefore did not change our procedure.

## Footnotes

1 https://support.10xgenomics.com/single-cell-gene-expression/software/downloads/latest#loupe

2 https://support.10xgenomics.com/genome-exome/software/pipelines/latest/what-is-long-ranger

3 https://github.com/PacificBiosciences/pbmm2

